# Stromule geometry allows optimal spatial regulation of organelle interactions in the quasi-2D cytoplasm

**DOI:** 10.1101/2023.04.27.538610

**Authors:** Jessica Lee Erickson, Jennifer Prautsch, Frisine Reynvoet, Frederik Niemeyer, Gerd Hause, Iain G. Johnston, Martin Schattat

**Affiliations:** Biology, Plant Physiology, Martin-Luther-University Halle-Wittenberg, Halle, Germany; Department of Biochemistry of Plant Interactions, Leibniz Institute for Plant Biochemistry, Halle, Germany; Department of Mathematics, University of Bergen, Bergen, Norway; Computational Biology Unit, University of Bergen, Bergen, Norway

**Keywords:** plastids, stromules, biotic stress, organelle interactions, optimal structures, Arabidopsis, Nicotiana

## Abstract

In plant cells, plastids form elongated extensions called stromules, the regulation and purposes of which remain unclear. Here we quantitatively explore how different stromule structures serve to enhance the ability of a plastid to interact with other organelles: increasing the effective space for interaction and biomolecular exchange between organelles. Strikingly, electron microscopy and confocal imaging showed that the cytoplasm in *Arabidopsis thaliana* and *Nicotiana benthamiana* epidermal cells is extremely thin (around 100 nm in regions without organelles), meaning that inter-organelle interactions effectively take place in 2D. We combine these imaging modalities with mathematical modelling and new *in planta* experiments to demonstrate how different the elongation of stromules (single or multiple, linear or branching) could be employed to optimise different aspects of inter-organelle interaction capacity in this 2D space. Stromule formation and branching is shown to provide a proportionally higher benefit to interaction capacity in 2D than in 3D. Additionally we find this benefit depends on optimal plastid spacing. We hypothesize that cells can promote the formation of different stromule architectures in the quasi-2D cytoplasm to optimise their interaction interface to meet specific requirements. These results provide new insight into the mechanisms underlying the transition from low to high stromule numbers during biotic stress, the consequences for interaction with smaller organelles, how plastid access and plastid to nucleus signalling is balanced, as well as the impact of plastid density on organelle interaction.

## Introduction

The process of enclosing the cellular lumen within membrane-bound compartments (compartmentalization) was crucial for cellular evolution and has led to a sophisticated and dynamic internal organisation found in today’s modern eukaryotic cells (Galdon et al., 2015; Johnston, 2019). These membrane-bound compartments, or organelles, enable the coexistence of multiple biochemical environments within a single cell. As a result, biochemical reactions can run under optimal conditions and even opposing reactions can exist in a single cell. However, as a consequence of compartmentalization many biochemical pathways are sequestered into two or more organelles (Lunn et al., 2007). One example for such a sequestered pathway in plant cells is photorespiration, which involves reactions in plastids, peroxisomes and mitochondria (Eisenhut et al., 2019).

One consequence of creating membrane bound organelles is that the sequestered compartments unavoidably obtain a shape, which is defined by their surrounding membranes. Interestingly, organelles often diverge from a simple spherical shape and undergo controlled morphological changes in response to external stressors or specific developmental processes (Mathur 2021; Fenton et al., 2021; Sheahan et al., 2004, 2005). It has been found that the shape of a membrane bound compartment can impact its biochemical performance (Lizana et al., 2008) and that the location and dynamics of organelles influences their function and capacity for exchange and interaction (Chustecki et al., 2021; Giannakis et al., 2022). This suggests that the shape of organelles is not random, but instead affords control over organelle function (Lizana et al., 2008, 2009).

An essential and ubiquitous instance of this structure-function relationship is found in plastids, organelles central for bioenergetics and metabolism in phototrophs. Plastids can exist in multiple specialised forms (Sierra et al., 2023), of which the chloroplast is among others responsible for light driven carbon fixation (Choi et al., 2021). Plastids are not only characterised by their biochemical flexibility and involvement in multiple biochemical pathways (Sierra et al., 2023; Bruncard et al., 2018; Weber et al., 2009), but are also known for their ability to change the shape of their enclosing membranes in drastic ways, by forming either irregular shapes or long tubules (reviewed in Gray et al., 2001 and Kwok et al., 2004). These thin, tubular extensions of the envelope membranes that are filled with stroma have been named stromules (Köhler et al., 2000). Stromules can vary in length, ranging from a few to over 65 µm, and have a thickness of about 0.4-0.8 µm. A notable characteristic of stromules is their dynamic nature, as they can rapidly extend, branch, kink, and retract within just a few seconds or minutes (Gunning et al., 2004). Despite being first observed over a century ago, stromules were largely ignored for years, as they are not easy to visualise under a light microscope (reviewed in Gray et al., 2001). However, with the development of fluorescence microscopy and the use of fluorescence proteins targeted to the stroma, stromules are now readily visible, and their prevalence throughout the *Viridiplantae* has been confirmed (Shaw et al., 2011, Gray et al., 2011, Delfosse et al., 2016, Mathur 2021).

Despite their evolutionary conservation and controlled formation during stress and development (reviewed in Mathur 2021), stromule function is still not fully understood. Inspired by microscopic observations in different tissues, a number of functions have been proposed, since their rediscovery in algae, in 1994 (Menzel et al., 1994), and in vascular plants, in 1997 (Köhler et al., 1997). Different authors reported that peroxisomes as well as mitochondria come into close proximity with stromules, inspiring the hypothesis that stromules might increase the interactive surface of plastids, facilitating a more efficient exchange of metabolites (Kwok et al., 2004 and 2004b, Schattat et al., 2011, Barton et al., 2017) for example during photorespiration or in between-organelle stress-induced transmission of defense signals (Serrano et al., 2016) and modulation of reactive oxygen species (Pastor et al., 2013; Arnaud et al., 2017). Further, observations describing stromules oriented towards nuclei, the cell periphery or plasmodesmata, led authors to speculate that stromules connect distant parts of the cell with the plastid body without the need for plastid repositioning (Kwok et al., 2004, Caplan et al., 2015). However, why some plastids extend stromules that are linear, some branched and some extend multiple stromules is not immediately addressed by this idea. Nor is it clear why plastids use stromule extension rather than motion to interact with distal partners.

Hypotheses on stromule function remain somewhat speculative, as the lack of stromule-specific mutants limits the capacity for direct experimental testing. Here, we propose an interdisciplinary approach combining detailed microscopic measurement of stromules, plastids, and their cytoplasmic environment, with mathematical and simulation modelling to evaluate how stromule formation impacts the interaction of plastids with their surroundings. We aim to use geometrical models of plastid and stromule structure, combined with optimisation approaches, to explore how the outer structure of plastids may be controlled to facilitate optimal interactions between cellular components.

## Results

### The cytoplasm is extremely thin in epidermal cells, limiting organelle interactions in the quasi-2D region

#### Stromule extension might increase the interactive surface of plastids in restricted geometries

**-** In previous work imaging organelle dynamics and interactions (Chustecki et al., 2021; 2022) it was observed that the vacuole in *Arabidopsis* hypocotyl epidermal cells acts to compress the cytoplasm into a thin layer adjacent to the cell wall, so that (for example) mitochondrial motion is often constrained to be largely planar (Chustecki et al., 2021). We reasoned that if this planarity restricts organelle motion and dimensionality, stromule extension and structure may facilitate more efficient exchange with different cellular regions than plastid motion. We therefore set out to identify the physical properties of the planar cytoplasm, and plastids embedded within it. We focus on epidermal cells as a model system that admit straightforward live imaging, without the high packing of organelles found in other tissues, but where stromules are known to form and interact with other compartments in response to different stimuli (Kwok et al., 2004b; Burton et al., 2017, Schattat et al., 2011b).

#### The cortical cytoplasm is a quasi-2D plane

In order to understand these spatial constraints, and how plastids and other organelles are embedded in the cytoplasm of the cell, we performed transmission electron microscopy (TEM) and confocal laser scanning microscopy on pavement cells in the leaf epidermis of wildtype *N. benthamiana* and *A. thaliana* plants, as well as on transgenic *N. benthamiana* cells. The images shown in Fig. 1 demonstrate that, as in the hypocotyl, these cells also host a large central vacuole that presses the cytoplasm with its content and the plasma membrane against the inner side of the cell wall, creating a very thin layer of cytoplasm (Fig 1A, Bb’’). As a consequence the largest portion of the cell lumen is occupied by the central vacuole (Fig. 1A, B and Bb’’). All other compartments reside within the thin cytoplasmic layer (Fig. 1C-F). As illustrated by transmission electron microscopy and confocal laser scanning fluorescence microscopy, many membrane bound compartments are wider than the cytoplasmic layer, e.g. plastids (Fig. 1Bb’ and C), mitochondria (Fig. 1D) and dictyosomes (Fig. 1E). As a consequence those compartments expose only a small part of their surface towards the cytoplasmic layer (Fig. 1 C-F), which represents the part of the compartments surface, which is the easiest to access for compartment-compartment interactions (Fig. 2A and B). For the large plastids the surface they expose to the cortical cytoplasmic layer is, relative to the rest of their surface area, especially small. Thus, for the interaction of a mitochondrion with a plastid only the small area of the plastid surface, which is exposed to the cytoplasmic layer, is available as depicted in the image shown in Fig. 1F. Only in a few areas within an epidermis cell, cytoplasm ‘accumulates’ providing a thicker cytoplasmic layer in which organelles can ‘pile up’ on top of each other, making the interaction of organelles with all parts of the plastid surface possible. Such areas typically form at the end of epidermis pavement cell lobes (Fig. 1G) and around the nucleus, which resides in a ‘cytoplasmic pocket’ (Fig. 1B and 1G). Only in rare cases have we observed organelles such as mitochondria and peroxisomes ‘climbing’ the much larger plastid body, gaining access to the plastid membrane exposed to the vacuolar membrane. Taken together, these observations in epidermal pavement cells demonstrate that membrane-bound compartments are predominantly constrained to interact with the plastid body in the area defined by the thin cytoplasmic layer and not with their complete surface (Fig. 2A), in the quasi-2D “flatland” of the cytoplasm (Abbott, 2008). Therefore, we concluded that for modelling approaches the interactions of plastids with their environment can be projected into a 2D space and still be representative (Fig. 2C-D).

**Fig. 1.**
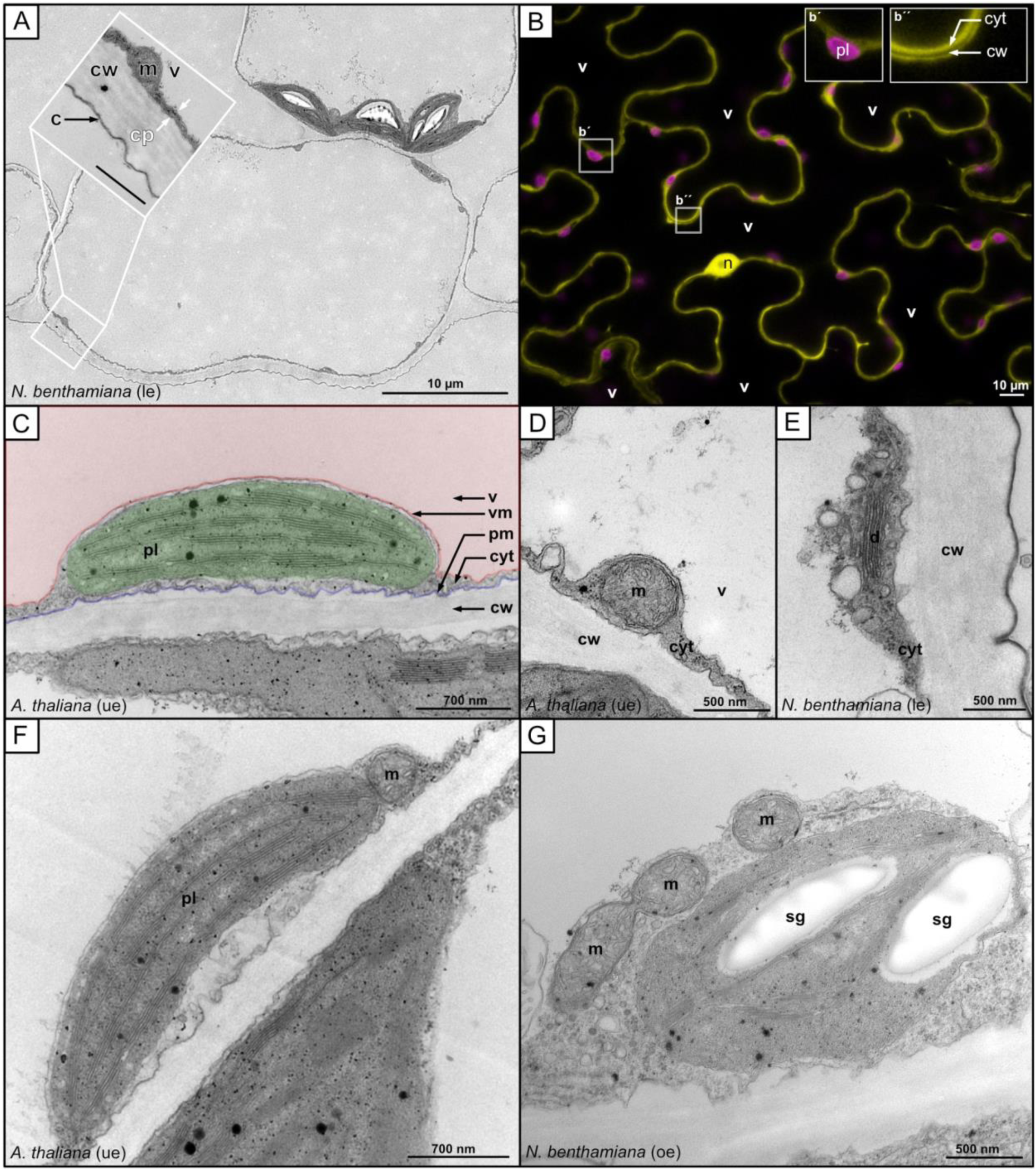
TEM and fluorescence images depicting organelles in the cortical cytoplasm. A) TEM image of a cross section through a lower epidermis leaf cell from *N. benthamiana*. The cytoplasm is visible as a thin dark gray lining along the inside of the lighter gray cell wall. Inset provides a detailed view of the cortical cytoplasm and the cell wall. B) Confocal laser scanning microscopy image of a lower leaf epidermal cell of *N. benthamiana* expressing untagged eGFP, which accumulates in the cytoplasm and the nucleoplasm. Inset b’ shows a chloroplast as highlighted by its chlorophyll-a auto fluorescence. Inset b’’ shows cytoplasmic GFP fluorescence of two cells separated by the non-fluorescent cell wall. C-G) TEM images of C) a plastid, D) a mitochondria, E) dictyosome as well as a plastid with F) one or G) three mitochondria in close proximity to it in an area of the cell with enriched cytoplasm. **v** = vacuole, **vm** = vacuole membrane, **pm** = plasma membrane, **cw** = cell wall, **c** = cuticle, **cyt** = cytoplasm, **m** = mitochondria, **pl** = plastid, **sg** = starch granule, **d** = dictyosome, **ue** = upper epidermis, **le** = lower epidermis.

**Fig. 2.**
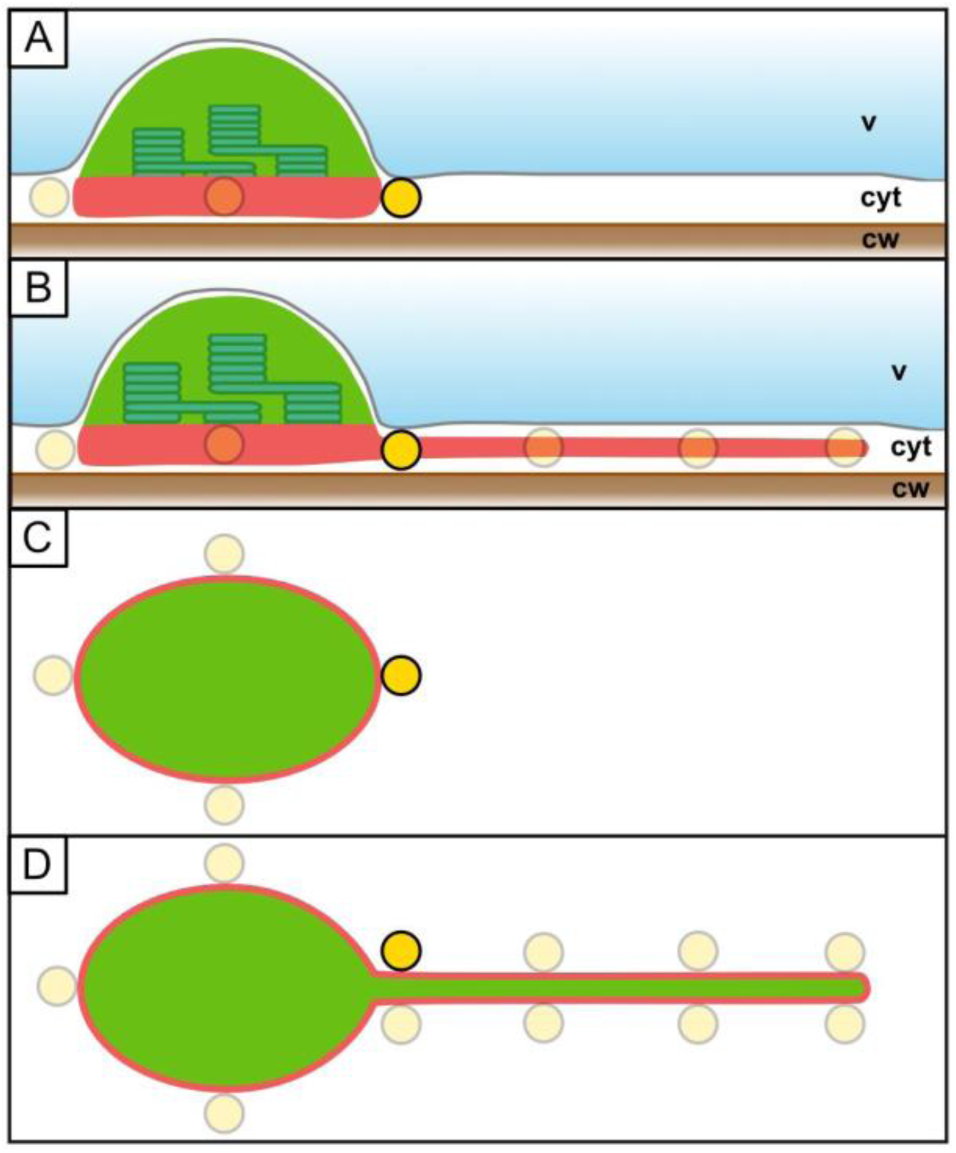
Spatial potential for plastid interactions in a quasi-2D cytoplasm. Plastids in the cortical cytoplasm (**cyt**) are pressed between the vacuole (**v**) and the cell wall (**cw**). **(A)** In the absence of a stromule extension, only a minor fraction of the plastid’s surface area is available for interactions with other organelles (yellow circle) residing in the cortical cytoplasmic layer (interactive surface area = red line and red shading). **(B)** Forming a stromule, reaching into the cytoplasmic layer, increases the exposed surface area, allowing for more interactions. **(C-D)** Due to the observed constellation of plastids, stromules and other organelles in the thin cortical cytoplasmic layer (see Fig. 1A-G), an epidermis pavement cell can be projected into a 2D space for modelling approaches and still represent the exposed interactive surface area of cortical plastids.

### Physical properties of stromules extending from flattened plastids into the thin layer of the cortical cytoplasm

Following this striking qualitative observation of a quasi-2D environment for organelle interactions, we next aimed to quantify the physical behaviour of plastids and the stromules extended from them within this quasi-2D plane. This data formed the foundation for later calculations.

#### Stromule formation can be induced by *Agrobacterium tumefaciens* inoculation

While stromule levels are relatively low under typical growth conditions (unchallenged), they are strongly induced when plants face biotic stress. In order to understand the benefit of stromules for a cell better it is of benefit to compare cells with basal stromule levels with those where stromules are induced. A suitable model system for such an approach is the treatment of *N. benthamiana* lower leaf epidermis with the *A. tumefaciens* laboratory strain GV3101(pMP90) (Fig. 3A and B). It is known from our previous work that stromule formation is triggered within 2-3 days post inoculation with higher optical densities of this strain (Erickson et al., 2014). In this case, stromules are specifically triggered by the activity of bacteria secreted trans-zeatin, which significantly alters the physiology of cells within the infiltration spot, leading for example to accumulation of soluble sugars and starch (Erickson et al., 2014). After two days, cells of inoculated leaf spots harbour a high number of stromules (over 50%) in contrast to untreated areas (below 10%) (Erickson et al., 2014). Therefore, we decided to obtain imaging data from unchallenged tissue as well as GV3101 (pMP90) treated tissue for quantitative plastid, stromule and cell analysis.

**Fig. 3.**
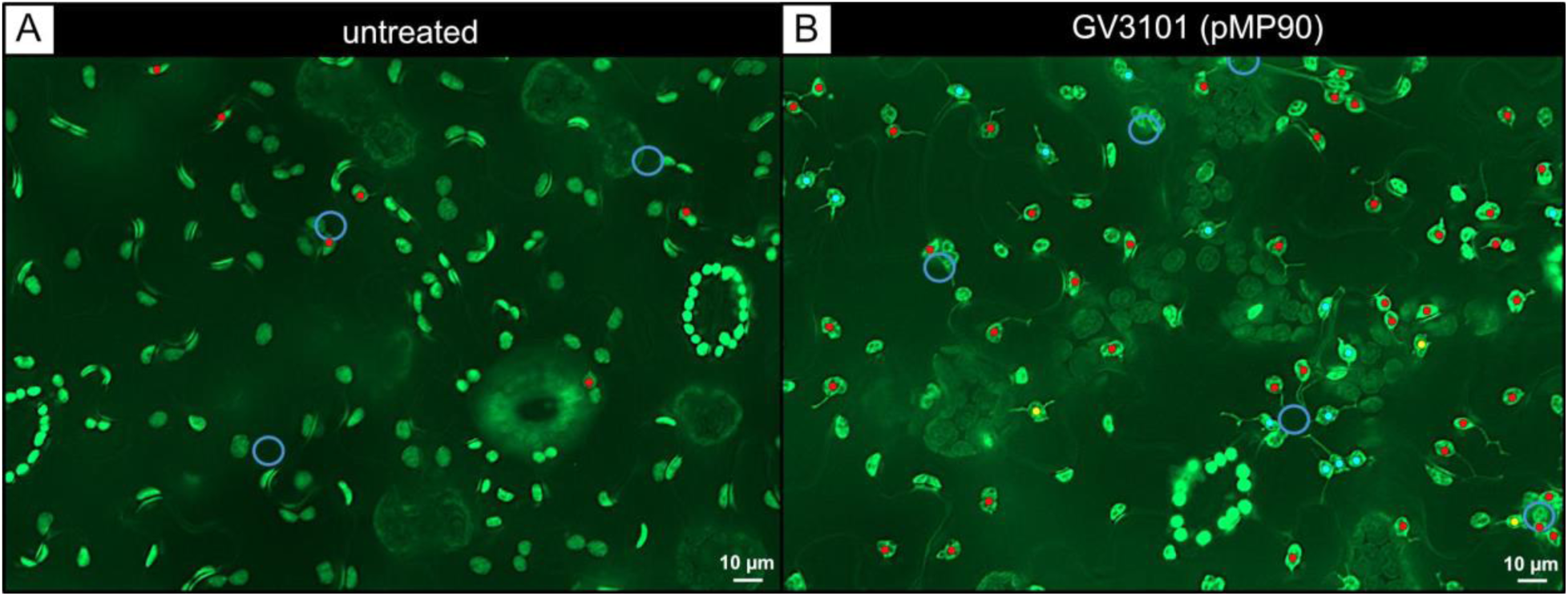
Extended depth of field fluorescence microscopy images of cells under control and stromule inducing conditions. Images show lower epidermis cells of *N. benthamiana* transgenic line FNR:eGFP#7-25 in untreated (A) and treated (B) leaf areas. A strong induction of stromule formation is visible 3 days post inoculation of *A. tumefaciens* GV3101 (pMP90) infiltration at a relatively high optical density (OD600nm = 0.8). blue circles = position of nuclei; dots = plastids with stromules, red = 1, cyan = 2, yellow = 3.

#### The cortical cytoplasm of the epidermis cells is around 100nm thick

Before we started to look at the different stromule characteristics we wanted to understand how thin the cytoplasm in our chosen model system (lower leaf epidermis cells of *N. benthamiana*) really is. Additionally we wanted to see if our assumption is true also for another model species and therefore included epidermis cells from *Arabidopsis thaliana* in our measurements. We quantified cytoplasm thickness from TEM images, finding that cytoplasm thickness in both *A. thaliana* upper leaf epidermis cells and *N. benthamiana* lower leaf epidermis cells was around 100 nm on average and was only very rarely observed to exceed 300 nm (Fig. 4A). Correspondingly TEM and Epifluorescence images of unchallenged *N. benthamiana* lower epidermis cells show clearly that plastids are pressed against the cell wall, appearing mostly oval shaped, when viewed from the side (pressed against the anticlinal cell wall) and appearing almost circular when viewed from the top (pressed against the periclinal cell wall; Fig. 1A, B, C, F and Fig. 3A and B). We further used the captured epifluorescence images to quantify the 3D structure of plastids in this environment, finding average values of 6.3 µm length, 5.0 µm width and 2.3 µm height (Fig. 4B).

**Fig. 4.**
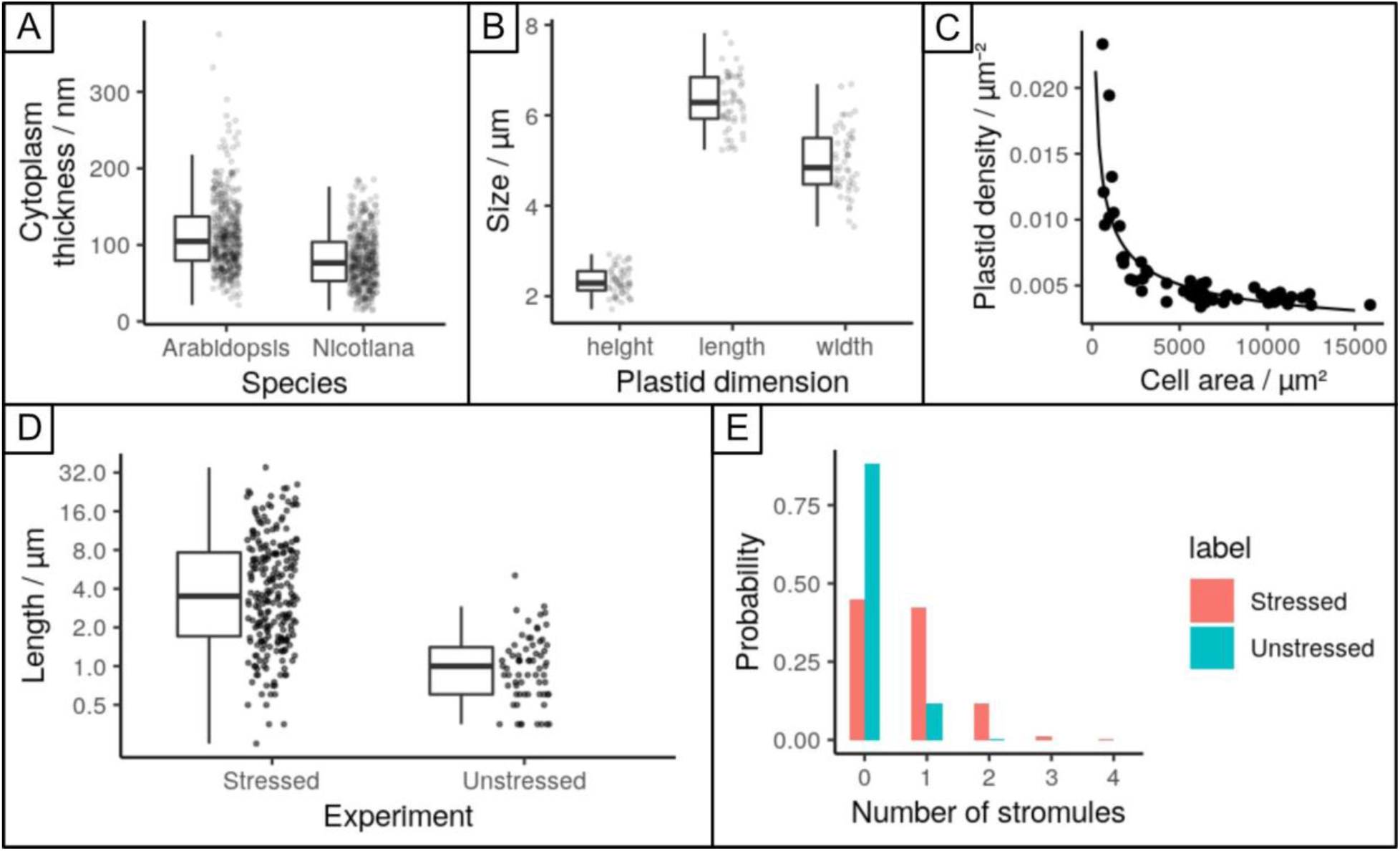
Quantitative measurements of cytoplasm, plastid, and stromule related statistics. (A) cytoplasmic thickness measured in TEM images of upper leaf epidermis cells of *A. thaliana* and lower leaf epidermis of *N. benthamiana*. (B) plastid dimensions length-width-height, measured in epifluorescence images. (C) Plastid density as measured in epifluorescence images as a function of cell size; curve gives a power-law fit. (D) Histogram of stromule length measurements in epifluorescence images (only unbranched stromules of plastids with a single stromule considered) in unstressed and stressed samples. (E) Proportion of plastids with different numbers of stromules in unstressed and stressed cases.

#### Plastid density in *N. benthamiana* is constant, but only in larger epidermis cells

We also measured the density of plastids in cells of different sizes (Fig. 4C). Strikingly, we found that plastids are more densely packed (up to around 0.014µm^-2^) in smaller cells, and that plastid density decreases drastically towards an asymptotic value (around 0.004µm^-2^) in larger cells. If these densities are interpreted as corresponding to the size of a “box” of cytoplasm in which only one plastid may be present, they correspond to characteristic plastid-plastid separations of between 8µm and 15µm. The behaviour of plastid density with cell size can be well fit (R^2^ = 0.8, Supplementary Fig. 1A) by a power law, though given the limited multiplicative range involved this should be interpreted cautiously [Stumpf & Porter, 2012]. In summary, the lower epidermis of *N. benthamiana* leafs is dominated by large cells with a rather low plastid density. In numbers this means that 8% of pavement cell area is occupied by small cells with plastid densities above 0.005 µm^-2^, the remaining 92% is covered by rather large cells with a constant low cell density below 0.005 µm^-2^.

#### In unchallenged and stressed cells the majority of stromules are short

Stromules are often found to vary substantially in length. To characterise this variability in our model systems we next measured the length distribution of stromules in unchallenged and stressed cells (Fig. 4D). We found that stromule length can be described in both conditions by a log-normal distribution (Supplementary Fig. 1B-C), with mean stromule length 5.8 µm and standard deviation of 5.8 µm; the log-normal model gives a 95% range of 0.53-25.3 µm. Stromule length clearly increases under stress, by a factor around 5-fold (Fig. 4D), with the mean of 5.8 µm compared to 1.2 µm in the unstressed case. Log-normal distributions are characteristic of quantities arising from independent multiplicative increments (like the scaling of individual elements), and are also observed in other dynamic, microscopic extension phenomena in biology, from the spines of neurons (Loewenstein et al., 2011) to cytoskeletal strands (Polizzi et al. 2021; Husainy et al.. 2010). In all these cases, including stromules, populations are dominated in numbers by short structures and longer structures are increasingly rare. While not a focus of this investigation, the mechanistic basis for this distribution could readily be, for example, the ongoing extension of a stromule by independent events where a motor protein attaches and elongates the stromule, scaling the length by a multiplicative factor before detaching. This is in agreement with current models for stromule elongation in which motor proteins are assumed to attach to the plastid surface, pulling out stromules by progressive movement along the respective cytoskeletal element.

#### Stromule count distributions per plastid are consistent with independent stromule initiations

**-** Fig. 4E shows the distributions of stromule count per plastid in stressed and unstressed conditions. This behaviour can be well fit by a Poisson distribution (Supplementary Fig. 1D-E), which is the expected behaviour if plastid initiation events are independent. In other words, the presence of one stromule on a plastid does not influence the probability that another is extruded; extrusion occurs independently and randomly with a characteristic rate. This rate is, on average, 0.69 per plastid in stressed samples (Fig. 4E). The stromule count per plastid also increases markedly under stress. Across samples, the Poisson model has a range of mean values 0.57-0.88 per plastid in the stressed case and 0.09-0.18 (mean 0.12) per plastid in the unstressed case, corresponding to the rate of stromule initiation increasing by between 3- and 6-fold under stress.

### Increases of physical capacity for plastid interaction due to stromule geometry

With the collected data describing different parameters of stromules, plastids and cells, we next asked if mathematical modelling could test the impact that stromules and plastid distribution have on plastid interactions with their cellular surroundings. For this we introduced two parameters describing two aspects of how plastids might interact with the cytoplasm and other organelles.

#### ‘Interaction region’ and ‘plastid access’ describe a plastid’s spatial capacity for interactions

**-** We define the “interaction region” of a plastid as the area of the planar region within a given distance *d* of the plastid’s boundary. We also define the “plastid access” of a given cellular point as the extent to which that point is surrounded by plastid: specifically, the proportion of the circumference of a circle of radius *d* drawn around the point that contains part of the plastid. The interaction region then describes how much of the quasi-2D cytoplasm is “available” to a plastid for interactions. The plastid access at a point describes how much of the circumferential surface of a putative organelle at that point could interact with the plastid.

#### Stromule extension increases the plastid’s interaction region and plastid access

We use a combination of mathematical modelling and simulation (Methods; Supplementary Text; Fig. 5) to describe how these quantities change with stromule length and geometry. We consider the “default” case of a plastid without stromules and several different arrangements of stromules (preserving total amount of membrane): a single long stromule, two shorter single stromules, and a branched stromule (Fig. 5A; Supplementary Fig. 2). Keeping the total amount of plastid membrane constant in both cases, stromule extension substantially increases the plastid’s interaction region across different proximities *d* (Fig 5A, E; Supplementary Text). Notable, the scale of this increase in interaction region is much more pronounced in 2D than would be the case in 3D (Supplementary Text; Supplementary Figs. 3-4B) – so the relative advantage of stromules for facilitating interaction is particularly high in the quasi-2D epidermal cytoplasm. A similar increase is found if the total plastid volume, rather than membrane, is kept constant (Supplementary Text; Supplementary Figs. 3-4A). In tandem, the extension of a stromule introduces some regions of the cell (adjacent to the point where the plastid is extruded) that have high plastid access (Fig 5B, F). A mitochondrion, for example, positioned at such a point will have part of its boundary adjacent to the main body of the plastid and another part of its boundary adjacent to the stromule, increasing the proportion of its boundary adjacent to the plastid (Fig. 5C).

**Fig. 5.**
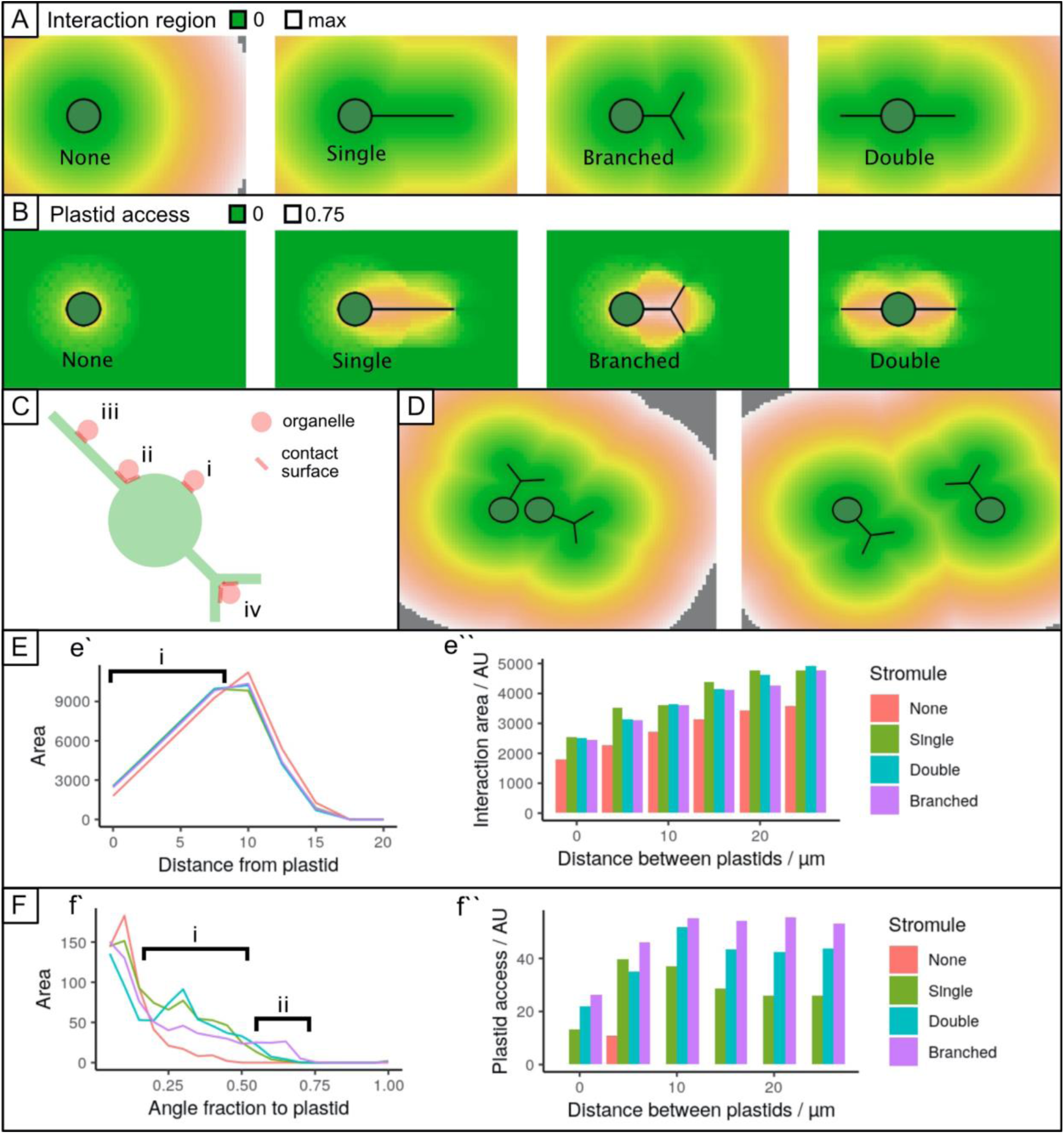
Interaction region and plastid access for different stromule geometries. (A) Interaction region of a plastid, plotted as the distance from a given point to the nearest part of the plastid. Distances range from 0 (within plastid) to L, the length scale of the model cell boundary. (B) Plastid access. A circle of radius d is drawn around each point. The proportion of the circumference for which a part of the plastid falls inside the circle is recorded. (C) Different arrangements of organelles and stromules. (i) Limited plastid access on the circumference of a circular plastid. (ii) Greater plastid access at the junction of a stromule. (iii) Limited axis along a stromule. (iv) High access at the branching point of a stromule. (D) Example of interaction region for a pair of plastids, changing with inter-plastid separation (left, 5um; right, 20um). (E) Distribution of cellular distances to the plastid for different stromule geometries (summary of (A)). (è) In region (i), stromule branching supports a greater area of closer cytoplasm than for the case without stromules. The behaviour for plastid pairs with different separations (summary of (C)) is shown on the right. (F) Distribution of plastid access through the cell for different stromule geometries (summary of (B)). (f’’) In region (i), stromule extension increases the amount of cytoplasm with high plastid access. In region (ii), branched stromules create some regions with very high plastid access. The behaviour for plastid pairs with different separations is shown on the right (e’’ and f’’’).

#### Only well-spaced plastids gain full benefit of stromule formation

We used this simulation approach to calculate the changes in these two quantities when two plastids, separated by a given distance, were considered together (Fig. 5D-F). For plastids separated by 5µm or less, comparatively little advantage is gained in either interaction region or plastid access. However, for higher separations, more substantial amplification occurs in both quantities, so that the advantage of having two plastids separated by a distance of 10-15µm is substantially greater. This length scale corresponds to the characteristic separation observed from the density measurements in Fig. 3C. The relatively higher advantage in a 2D geometry also holds here (Supplementary Text), highlighting the value of stromule extension for multiple plastids in the quasi-2D environment.

### Stromule branching increases the accessibility of a plastid for other cellular regions

#### Branched stromules increase the plastid access of cytoplasmic regions

From the perspective of interaction region alone, assuming a constant amount of membrane, branching is always at least slightly worse than extending a single long stromule (Fig 5B, F). This is because the interaction region is “double counted” in the region of a branch point: extending a long stromule will reach more cytoplasm than branching in the vicinity of an existing stromule. However, branched stromules achieve a different geometrical goal, which is increasing the plastid access of different cellular regions. A point near the junction of a branched stromule is more surrounded by plastid than one near a single continuous branch (Fig. 5C). A mitochondrion positioned at the branch point, for example, would have a majority of its boundary within a small distance of the plastid, whereas one adjacent to a single stromule would have a limited amount of its boundary adjacent to the plastid. Correspondingly, branched stromules create regions of very high plastid access, where an organelle is mostly surrounded by plastid (Fig. 5B, F). To maximise interaction region with a branch point, theory predicts that branching angles should be as close to 120° as possible (Supplementary Text; Supplementary Fig. 5). This agrees with observations of branching angles between 100°-140°, centred on 120° (Schattat et al., 2011b).

### Observed stromule behaviour under stress enhances inter-organelle interaction capacity

Taken together, these theoretical and simulation results show that a tradeoff exists for stromule structure – mirroring the “physical-social” tradeoffs observed in other organelle interaction behaviour (Chustecki et al., 2021). Single linear stromules maximise the region of the cell close to the plastid. But stromule branching allows other organelles to maximise the amount of their perimeter in contact with the plastid. Branching at 120° provides a resolution to this tradeoff, where the loss of proximal region is minimised while branching is supported. This theory then predicts that, depending on the relative weighting of two competing cellular priorities (maximising a plastid’s capacity for interaction, and maximising an organelle’s capacity to interact with the plastid), stromule structure should favour either linear or branching at 120°. These are the structures which are overwhelmingly most commonly observed in cells (Fig. 3; Schattat et al., 2011b), suggesting that biology adopts these optimal resolutions to the challenge of regulating organelle interactions in quasi-2D. At the same time, multiple plastids separated by distances over 5µm amplify the advantages from these structures. Once again, the characteristic separation observed in cells (8-15µm) fulfils this condition. Taken together, these observations support a picture where stress induces a response in stromule structure that supports increased inter-organelle communication. The dramatic increase in stromule initiation and length that we and others observe serves to increase the effective interaction area of a plastid. This effect is particularly pronounced in the quasi-2D cytoplasm in epidermal cells. The extent of stromule branching controls a tradeoff between access of the plastid to other cellular components (our “interaction region”) and access of other organelles to the plastid (our “plastid access”).

## Discussion

The impact of organelle shape on organelle function and on cell viability is difficult to address experimentally. This is mainly due to the absence of morphology-specific mutants, especially in the case of plastid stromules. Thus, alternative approaches to evaluate the importance of organelle morphology are a promising avenue of enquiry. Here we have combined electron microscopy and fluorescence imaging to reveal the physical details of organelle interaction in the cytoplasm with a focus on plastids, showing that the physical space in which such colocalisations must occur is thin to the point of being a quasi-2D “flatland” (Abbott, 2008). Given this restricted space, extension of (quasi-1D) stromules powerfully increases the proportion of the cell with which the plastid can interact: mathematical modelling demonstrates the relative amplitude of this effect in 2D versus less restricted 3D space. Branched stromules do not serve to increase this interaction region, but instead create regions of the cytoplasm more surrounded by plastid content, so that organelles positioned there have amplified access to the plastid through their membrane. We describe how the observed stromule structures in plant cells constitute a range of optimal solutions to a tradeoff between maximising a plastid’s capacity for interaction, and maximising an organelle’s capacity to interact with the plastid. In the following paragraphs we discuss more specifically how the obtained results can help to gain new insight into observed and not fully understood stromule behaviour and how this will guide future research directions.

### Insights into the mechanism of stromule initiation

Although in the past years fundamental progress has been made to understand how individual stromules are established and shaped by the action of actin and microtubule dependent processes (Erickson et al., 2018; Kumar et al., 2018; Kwok et al., 2003), it is still not known how the transition from a low stromule frequency state to an induced stromule state is achieved for a complete plastid population of a cell. Among the open questions are: How is the decision made which plastid will form a stromule? How is it decided if a plastid forms one or more stromules? How is the decision made to form a single straight stromule or if a stromule branch is initiated? Knowing the answer to these questions would provide important insights into how the plant cell integrates stress signals to mount a complex morphological response. Our analysis starts to answer some of these questions. The Poisson distribution of stromule initiation events among plastids (Fig. 4E) suggests that this process is random and independent in *N. benthamiana* lower epidermis cells, and that the possession of one stromule does not increase the likelihood of gaining a second. In other words, the extension of stromules is unlikely to occur via the concentrated, local activation of the required stromule building ‘machinery’, but rather evenly distributed and activated cellular components are expected to contribute. So far, this fits with previous observations that stromule extension in *N. benthamiana* relies on microtubules, which are found throughout the cell cortex (Erickson et al., 2018; Kumar et al., 2018). Based on this we further speculate that the machinery responsible for a low stromule frequency state is similar to the one active when the cell transitions to a high stromule state and only its activity is increased. Behaviour in *A. thaliana* upper epidermis pavement cells indicates that, in some tissues, stromule forming activity might be focused in specialised areas of the cells, such as in close proximity to the nucleus. Our current dataset misses this information, so it is up to future experiments to address if a positional effect on stromule formation in *N. benthamiana* exists. However, the random nature of stromule initiation events observed in *N. benthamiana* epidermis is compatible with a picture branches may represent stromules that have independently initiated from a previously formed stromule rather than the plastid body, implying that for forming a new tubule the plastid membrane has no priority over the plastid body. If true, this would emphasise that the machenry at work to form stromules and branches is likely the same for both processes. Also here future work is needed to obtain more specific data, which will allow us to address this in more detail.

### The consequences of increased plastid access

Physical membrane-membrane contact between organelles is thought to increase the exchange of signalling molecules, lipids and, in some cases, may also contribute to organelle positioning (Baillie et al., 2020). Our modelling of membrane accessibility tells us something about how interactions with smaller organelles might occur. The rounded surface of the plastid body not only has limited contact surface, but movement by the smaller organelle in almost any direction will take it away from the plastid. This is in stark contrast to a plastid with a stromule, which has more contact surface, and since the smaller organelle is surrounded on several sides (especially in the case of a branched stromule), the likelihood of the smaller organelle losing contact with the plastid is less. Future considerations include how an increase in access might influence other factors, such as the duration of interactions in the dynamic cellular environment. For example, peroxisomes and mitochondria seem to ‘pile up’ against stromules when caught up in the cytoplasmic stream (Kwok and Hanson, 2004; Gunning, 2005). ‘Catching’ of smaller organelles in these ‘high access’ zones created by stromules could represent one means by which stromules not only increase contact area, but also increase the duration of interactions in a dynamic cellular environment. Here, detailed time lapse imaging of plastids together with other organelles will help to form the foundation for modelling in this direction.

### Balancing nuclear and cytosolic interactions of plastids

It is well known that plastid positioning is not random, but tightly controlled through associations with actin (Oikawa et al., 2008). Fluorescence images captured for this study show that plastids in epidermis pavement cells of unchallenged *N. benthamiana* leaves are quite evenly distributed within the cytosol (Fig. 3A and B). In addition, stromule frequency is minimal in the absence of stress (Fig. 1A). It is readily accepted that when plants are confronted with stress stimuli, interactions and communication between organelles, cytosol and plasma membrane increases in importance (Medina-Puche and Lozano-Duran, 2022). Recent attention has been paid to instances of plastid repositioning to the nucleus, which readily occurs upon confrontation with biotic stress, such as exposure to pathogen-derived effectors and peptides that trigger immune responses (Erickson et al., 2014, 2018; Caplan et al., 2008, 2015; Ding et al., 2019; Prautsch et al., 2023). In this context, plastid proximity to the nucleus is cited as important for the efficient transfer of retrograde signals such as H_2_O_2_ and proteins from plastids to nuclei (Caplan et al., 2008, 2015; Park et al., 2018; Ding et al., 2019). One would expect that, if plastid relocation to the nucleus is essential for responding to stress, that most plastids would be found at the nucleus, especially during the integration of strong stress signals, such as those associated with ETI. However, at least in the context of our work, inoculation with GV3101 (pMP90) (Fig. 3A and B), and triggering of ETI by the expression of XopQ results in only a subset of plastids relocating to the nucleus (Erickson et al., 2014; Prautsch et al., 2023), while the rest remain evenly spread (Fig. 3B and Supplementary Fig. S6). The results we obtained in our modelling provide a possible explanation for this contradiction. While clustering of plastids at the nucleus decreases the diffusion distance for retrograde signals, this arrangement sacrifices plastid interaction area. The fact that the even spacing of plastids is largely maintained during stress responses implies that cytosolic interactions, other than those with the nucleus, are also of importance during biotic stress. We calculated that the greatest gains in overall interaction region actually occur when evenly spaced plastids extend stromules, which is what we observed in our experiments. It would seem that a balance between plastid repositioning and stromule extension (total access of cytoplasm) is struck to mount the appropriate response to a given external stimulus.

### Are stromules obsolete in densely packed mesophyll cells?

Emitting a stromule is not the only mode by which plastid access increases. Although we have only modelled a scenario with two plastids, we have already shown that plastid access increases, even in the absence of a stromule, but only when plastids are close enough together (Fig. 5). This benefit gets lost again when plastid distance increases. In this case organelles caught between two plastids gain access to both plastids at the same time. So although we have found that stromules have a big impact on both interaction area and access in epidermal cells, which have low plastid densities, we would expect that mesophyll cells, which are densely packed with plastids, have higher average plastid access even without forming stromules, and that the impact of stromule formation would be reduced. Interestingly, mesophyll cells have been reported to have fewer stromules than epidermal tissues (Hanson et al., 2008), lending support for this idea. Experimental evidence is provided by a study conducted in *Nicotiana tabaccum* seedings, which showed that stromule length in hypocotyl cells, undergoing skotomorphogenesis, is negatively correlated with plastid density (Waters et al., 2004). This means that with increasing cell elongation and the resulting plastid density decrease, the authors observed a strongly correlated stromule length increase. Taken together, including more than two plastids and different plastid densities in our modelling approaches will allow us to gain a more holistic understanding on this aspect of plastid morphology in different cell types and developmental stages.

## Conclusion outlook

In order to overcome the current experimental limitations for addressing the impact of plastid shape and shape change on organelle function and cellular organisation, we combined measured morphological parameters for plastids and stromules with mathematical modelling and simulation. The obtained results allowed us to gain insight into the consequences, which stromule formation implies for plastid-cytoplasm interactions. The obtained results allowed us to look at open questions in this field from a new angle, lending new support for existing hypotheses and suggesting new explanations for so far unexplained observations. All together our study shows that this cross-disciplinary approach has potential to help form informed hypotheses and to guide future experimental design. Because we are here looking at the general consequences of shaping plastid membranes and plastid positioning our findings are of general nature across plastid-related processes involving interaction of the plastid with its surroundings. The future challenge will be to test if the predicted impact can be measured using physiological methods. Here the development of tools to manipulate organelle shape and positioning will be crucial. It will also be important to consider that stromule formation is likely the result of independent mechanisms (Erickson et al., 2018b), which might have a different morphological impact (Köhler et al., 2000; Waters et al., 2004).

## Material and Methods

### Plant material and growth

In this study, *Nicotiana benthamiana* wild-type and a transgenic plant line, highlighting the plastid stroma with eGFP (FNR:eGFP#7-25), was used (for details see Schattat et al., 2011b). Plants were cultivated in greenhouse chambers under long-day conditions, with 16 h of daylight (23°C), 8 h of darkness (19°C) and 55% relative humidity. *Arabidopsis thaliana* wild-type col-0 plants were cultivated in walk-in chambers under short-day conditions, with 8 hours of light (21°C) and 16 hours of darkness (17°C) and a relative humidity of 60%.

### Transmission electron microscopy

Leaf disks (3 mm) of *Arabidopsis thaliana* and *Nicotiana benthamiana* were fixed with 3% (w/v) glutaraldehyde (Sigma, Taufkirchen, Germany) in 0.1 M sodium cacodylate buffer (SCB; pH 7.2) for 3 h at room temperature. After fixation, the samples were rinsed in SCB and postfixed with 1% (w/v) osmium tetroxide (Carl Roth) in SCB for 1 h at room temperature. Subsequently, the leaf segments were rinsed with water, dehydrated in a graded ethanol series (10%, 30%, 50%, 70% (containing 1% (w/v) uranyl acetate), 70%, 90% and 2×100% for 30 min each), infiltrated with epoxy resin according to Spurr (1969), and polymerized at 70°C for 18 h. Ultrathin sections (70 nm) were made with an Ultracut E ultramicrotome (Leica, Wetzlar, Germany). Sections were transferred to Formvar-coated copper grids, post-stained with uranyl acetate and lead citrate with an EMSTAIN (Leica) and observed with a Libra 120 transmission electron microscope (Carl Zeiss Microscopy GmbH, Jena, Germany) operating at 120 kV. Images were taken by using a dual-speed on-axis SSCCD camera (BM-2k-120; TRS, Moorenweis, Germany).

### Infiltration of Agrobacteria into *N. benthamiana* leaves

Standard transient expression protocols were used to carry out the infiltrations. The bacterial ‘overnight’ culture was pelleted, re-suspended, and then left to incubate in infiltration media containing acetosyringone for approx. 2h at room temperature. The infiltration media, called AIM, consisted of 10 mM MgCl2, 5 mM MES pH 5.3, and 150 μM acetosyringone (Sigma-Aldrich, Deisenhofen, Germany). After incubation, the suspension’s optical density was adjusted to either OD600nm = 0.2 (*xopQ:mOrange2* expression) or OD600nm = 0.8 (*dsRed* GV3101 (pMP90) treatments). A needless syringe was used to infiltrate the *A. tumefaciens* strains into the intercostal fields of the lower side of the third or fourth leaf of 6-8 week old *N. benthamiana* plants. After 3 days post infiltration, leaf discs from the infiltrated areas and a non-infiltrated control were harvested, and the lower leaf epidermis was observed using either an epifluorescence or confocal microscope.

### Agrobacterium strains

For stromule induction two different *Agrobacterium tumefaciens* based inducers where used. A) Transient expression of the effector triggered immunity inducing effector Xanthomonas Outer Protein Q (XopQ; Prautsch et al., 2023) was mediated by *A. tumefaciens* strain GV3101 (pMP90). This strain carries resistances to rifampicin, gentamicin as well as kanamycin. B) Induction of stromules by exposure of epidermis cells to high concentrations of cytokinin was achieved as described in Erickson et al., 2014 by inoculating leafs with an OD600 = 0.8 of the trans-zeatin producing *A. tumefaciens* strain GV3101 (pMP90). In order to control for the presence of bacteria in the inoculated leaf areas the strain carried the T-DNA vector *pCP60:DsRed* mediating expression of the un-tagged fluorescent protein DsRed2. This strain carries resistances to rifampicin, gentamicin as well as kanamycin.

### Plasmids

The *xopQ-mOrange2* expression construct was described previously (Erickson et al., 2018). Expression of the XopQ-mORANGE2 fusion is driven by a *35S* promotor. The T-DNA vector *pCP60-35S-DsRed2* (untagged dsRed2) was kindly provided by C. Papst (lab of U. Bonas, MLU, Halle (Saale), Germany) and has been used previously for similar purposes (Erickson et al., 2018; Prautsch et al., 2023). For labeling the cytoplasm, untagged eGFP was expressed from pGGA2 (Schulze et al., 2012). pGGA2 harbours the pBGWFS7 backbone with spectinomycin resistance cassette, allowing for selection in *Agrobacterium*, with a 35S promoter and terminator.

### Confocal and Epifluorescence Imaging

Harvested leaf discs were vacuum-infiltrated with tap water right before imaging and mounted on glass slides. Subsequently z-stacks of the lower epidermis were collected. **Epifluorescence imaging:** For image acquisition, an epi-fluorescence microscope (AxioObserver Z1) setup from Zeiss (Jena, Germany) equipped with an X-Cite fluorescence light source and a MRm monochrome camera (Zeiss, Jena, Germany) was used. GFP fluorescence was recorded using a 38 HE filter cube (Carl Zeiss AG, Jena, Germany). mORANGE2 fluorescence was recorded utilizing the 43 HE filter cube (Carl Zeiss AG. Jena, Germany). The microscope manufacturer’s software (ZenBlue, Zeiss, Germany) controlled image acquisition. All images were captured using a 40x / 0.75 NA EC PLAN NEOFLUAR lens. **CLSM imaging:** Images used to calculate plastid density as well as the stromule reaction toward *XopQ:mOrange2* expression, a LSM 780 from Zeiss, Germany, was used (excitation: Argon laser line 488 nm, HeliumNeon laser line 633 nm; MBS 488 nm / 561 nm / 633 nm; emission detection: eGFP 499 nm - 606 nm, chlorophyll A 647 nm - 721 nm; lens: Plan-Apochromat 20x/0.8). Image used to demonstrate thickness of cytoplasm in Fig. 1B was captured with a Leica Stellaris CLSM (excitation: 474 nm; emission eGFP 489 nm - 537 nm, chlorophyll A 671 nm - 697 nm; lens: HC PL APO CS2 40x/1.10 WATER).

### Image processing and quantification of stromule/plastid parameters

#### Stromule and plastid parameters

For measuring stromule and plastid parameters images of infiltrations with OD600nm = 0.8 of GV3101 (pMP90) *pCP60:35S:DsRed* as well as images of untreated FNR:eGFP#7-25 plants were used. In order to obtain 2D extended depth of field images for quantification, single images of the z-series of each channel were first exported into separate file folders and subsequently combined into single images using software and procedures described in Schattat and Klösgen (2009). For the quantification of stromule frequencies (SF%), we measured the proportion of plastids with at least one stromule (Erickson et al., 2014) with the help of the MTBCellCounter (Franke et al., 2015). Stromule length was measured by tracing stromules with the line tool and using the ‘skeletonize (2D/3D)’ as well as the ‘analyze skeleton’ function of Fiji (Schindelin et al., 2012). Number of plastids with different numbers of stromules was counted manually with the help of the MTBCellCounter (Franke et al., 2015). Plastid proportions where measured with the ‘free hand line’ tool in images of not treated leaves using Fiji (Schindelin et al., 2012). Plastid density measurements were done in maximum intensity projections along the z-axis of 3D z-stacks. Plastid density is expressed as the number of plastids per area unit for each analyzed cell. In order to obtain plastid density, cells were outlined in stacked bright field images (Schattat and Klösgen, 2009). Outlines were used to measure the projected cell area in µm2, using Fiji (Schindelin et al., 2012). Superimposed outlines were used to count plastids in each outlined cell. Both measurements were used to calculate plastid density. **Cytoplasm thickness:** Cytoplasm thickness was measured in TEM images of ultrathin sections by adding ‘line’ drawing elements connecting the vascular membrane (tonoplast) with the plasma membrane, using Fiji (Schindelin et al., 2012). A random grid was used to choose measuring spots, avoiding measuring across organelles. Analog to stromule length measurements, generated lines were analysed for length in batch.

### Mathematical and computational models

Mathematical modeling was based on the representation of plastid bodies and stromules as simplified geometric objects like circles, rectangles, and cylinders, and calculating regions adjacent to these objects as a function of their arrangement (Supplementary Text). Simulations were carried out by embedding a set of line segments representing the plastid circumference and stromules of various geometries in the x-y plane. The plane was then partitioned into pixels. The distance of each pixel to its nearest line segment was computed to construct the interaction region of a plastid. Raytracing was used to compute the plastid access of a pixel. A ray was extended at angle *θ* from the pixel of interest and if the ray intersected a plastid line segment within a distance *D* of the pixel, an encounter at angle *θ* was recorded. θ was scanned from 0 to 2π radians in increments of 0.1, and the proportion of angles for which an encounter was reported is recorded.

Plastid bodies were assigned radii of 2µm. Line segments corresponding to stromules had conserved length *l* = 10µm, which could be partitioned as a single stromule of length *l*, two stromules of length *l*/2 extended from opposite sides of the plastid body, or a branched stromule where a segment of length *l*/3 branches to two other length *l*/3 segments all at 120° angles. The angle of extension of the first stromule was varied randomly in different simulation instances. We simulated the cases of a single plastid, and two plastids with their centres separated by horizontal (x) distances of 5 to 25µm in 5µm steps. Simulation was carried out using custom code written in C; this code and R implementations of the mathematical results, statistical analyses, and plotting scripts for all modelling is freely available at github.com/StochasticBiology/stromule-geometry.

## Acknowledgments

We thank Bianca Rosinsky as well as Siegfried Platzer for technical assistance with horticulture. We also thank Simone Fraas for the preparation and sectioning of the leaves for electron microscopy.

## Supplementary Information

### Statistics and model fits

Fig. 1 shows model fits for plastid density, stromule length, and stromule count distributions.

### Simulation output

Fig. 2 shows further examples of simulation output for different plastid separations and stromule geometries.

### Linear stromule extension

We picture a plastid as a sphere with initial radius *r*_0_. A cylindrical stromule of radius *ρ* and length *l* can be extended from the surface of the sphere. There are two possible pictures we can consider: (i) total plastid surface area is conserved (constant amount of membrane); (ii) total plastid volume is conserved (constant amount of stroma). In each case, as stromule length *l* increases, the radius of the spherical part of the plastid will decrease, as membrane/stroma is ‘donated’ to the stromule. We write *r* for the radius of the spherical part.

The total surface area and volume of sphere and stromule are

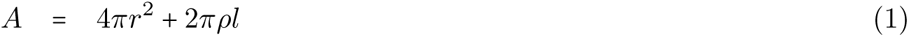

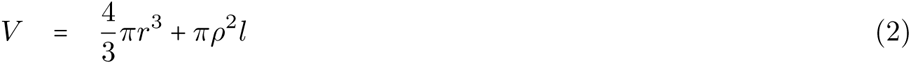

Under (i), total area is conserved, so 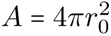 and

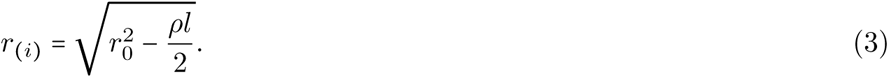

Under (ii), total volume is conserved, so 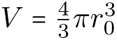 and

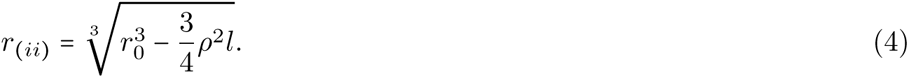

Reflecting the fact that sphere and stromule are embedded in a thin layer of cytosol, we will consider the cytosol as a quasi-2D plane (Fig. 3A-B). We are interested in 𝒜 (*d*), the area of the plane that lies within a distance *d* outside the plastid.

From the labelled areas in Fig. 3A-B,

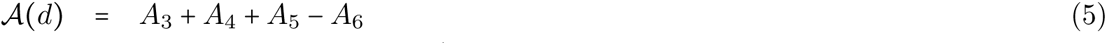

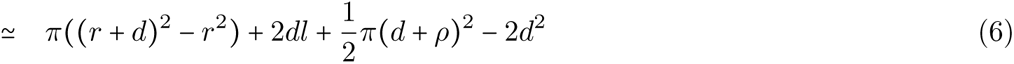

where some approximations have been made and the subtraction of *A*_6_ is to avoid double counting in that region. We can insert Eqns. 3-4 and simplify to obtain

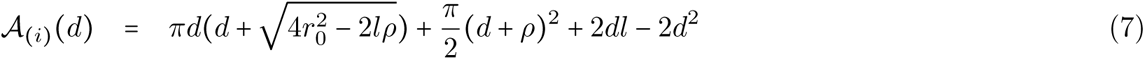

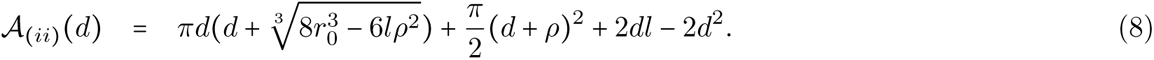

Taking some images from [?] as a guide, we estimate as characteristic values *r*_0_ = 2*µm* (plastid radius), *l* = 10*µm* (stromule length), *ρ* = 0.25*µm* (stromule radius). Then Fig. 4A demonstrates that even a moderate stromule length dramatically increases the area of the cytoplasmic plane within a given distance of the plastid.

**Figure 1:**
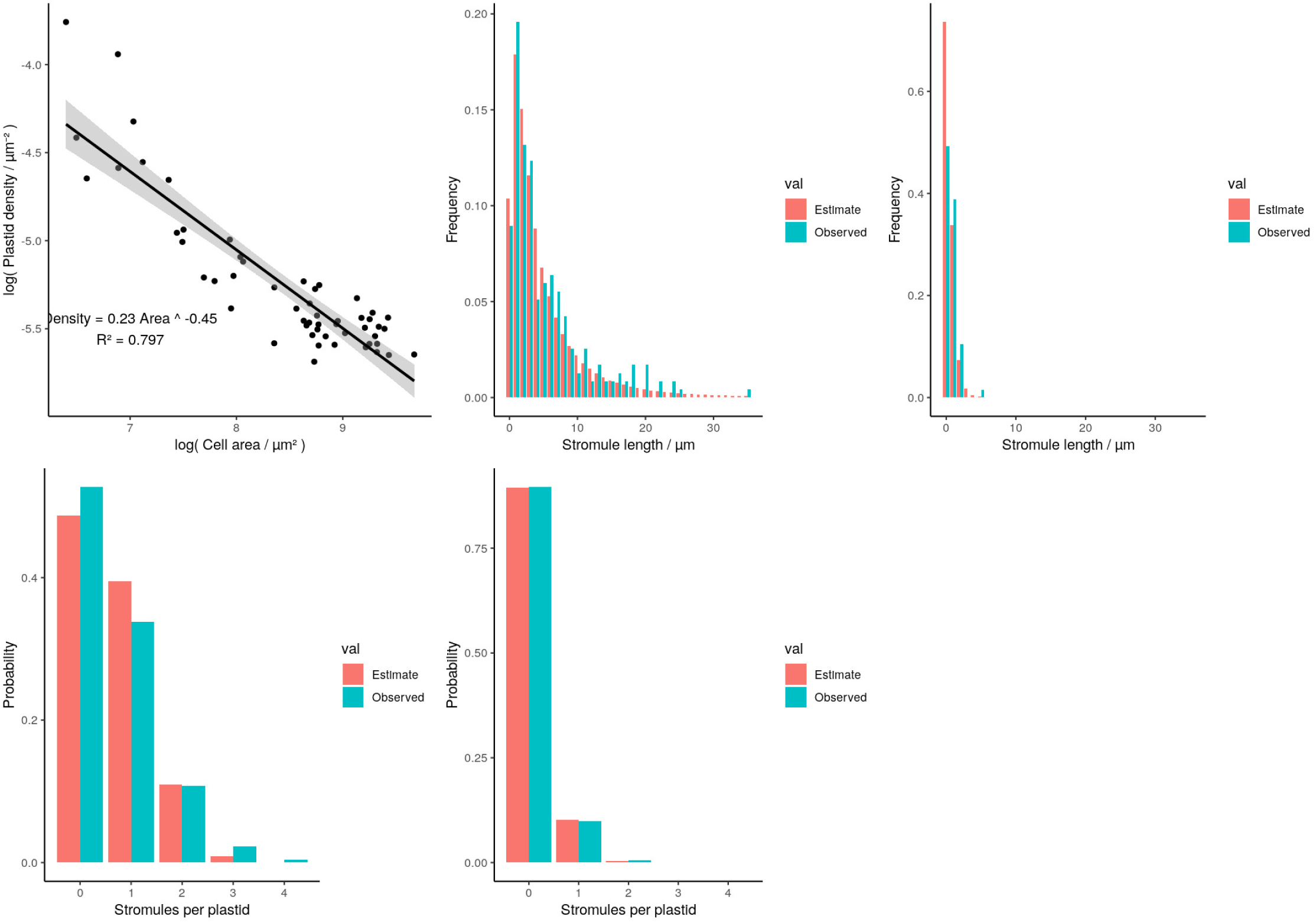
Model fits for plastid and stromule statistics. (A) Power-law fit relating plastid density to cell area. (B) Estimated log-normal distribution fitting observed stromule lengths in stressed samples. (C) Estimated log-normal distribution fitting observed stromule lengths in unstressed samples. (D) Estimated Poisson distribution fitting stromule counts per plastid in stressed samples. (E) Estimated Poisson distribution fitting stromule counts per plastid in an unstressed samples.

**Figure 2:**
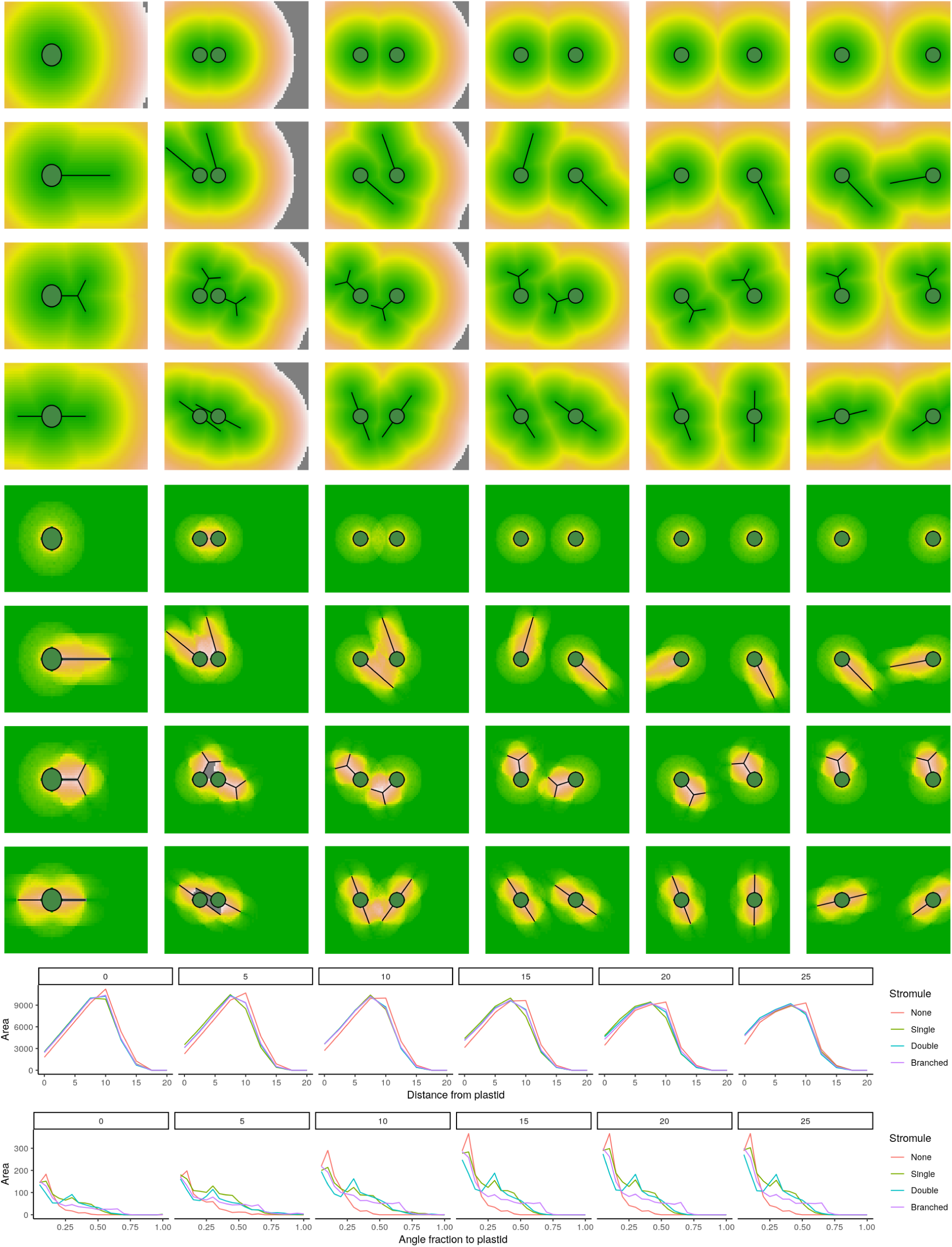
Simulation outputs. (top) Interaction region and (centre) plastid access for (rows) different stromule structures and (columns) single plastids, then pairs of plastids separates by 5, 10, 15, 20, 25*µ*m. (bottom) Interaction region and plastid access quantified from these simulations as in the main text; different panels give different plastid separations (0, single plastid).

**Figure 3:**
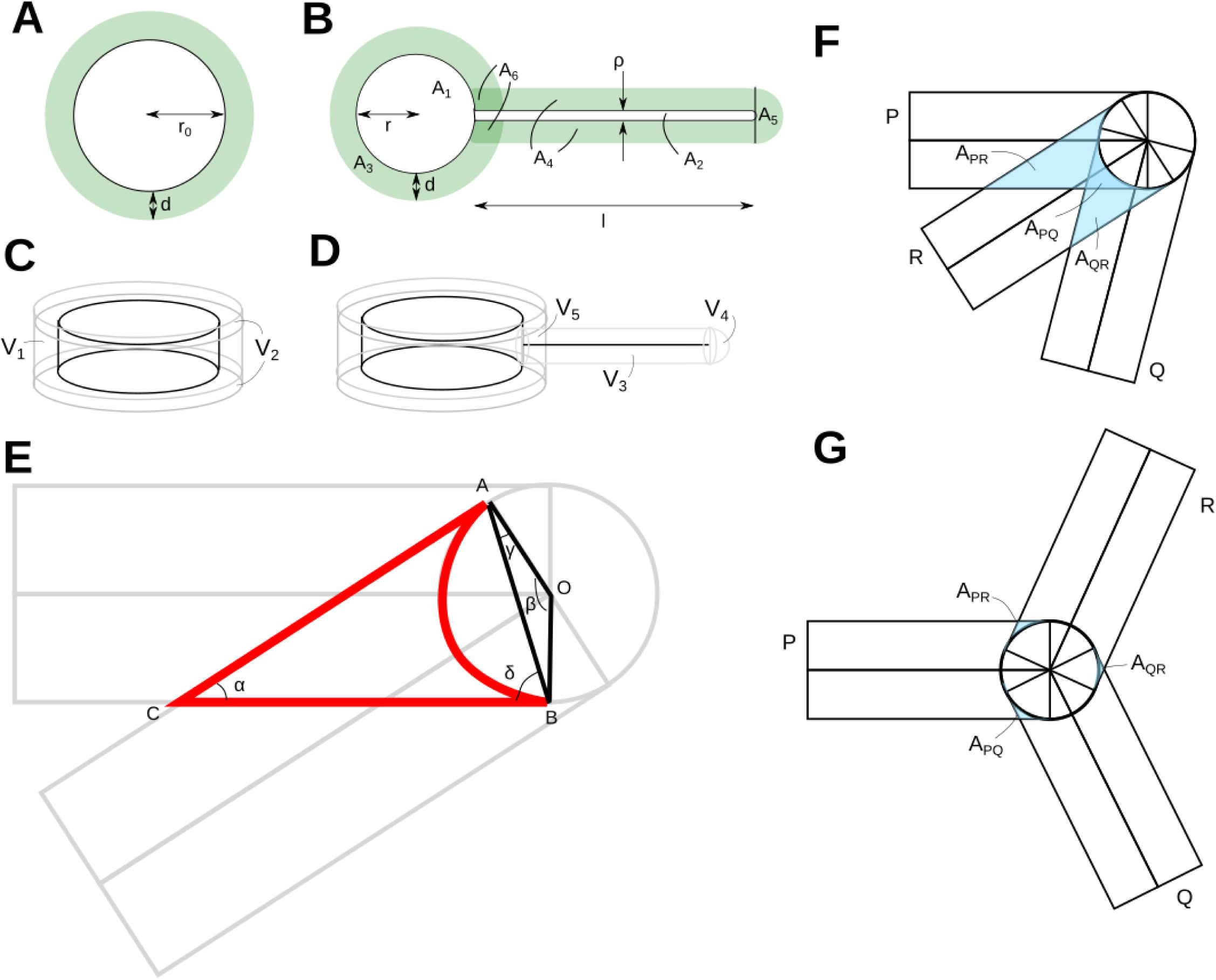
Geometric models for stromules. (A-D) Shells surrounding linear stromule extension model, in 2D with (A) no stromule and (B) extended stromule; and in 3D with (C) no stromule and (D) extended stromule. (E) Overlap region of shells around two stromule segments. (F) Overlapping regions around a branch point where one segment falls ‘between’ two others; (G) overlapping regions around a branch point where one segment falls ‘outside’ two others.

**Figure 4:**
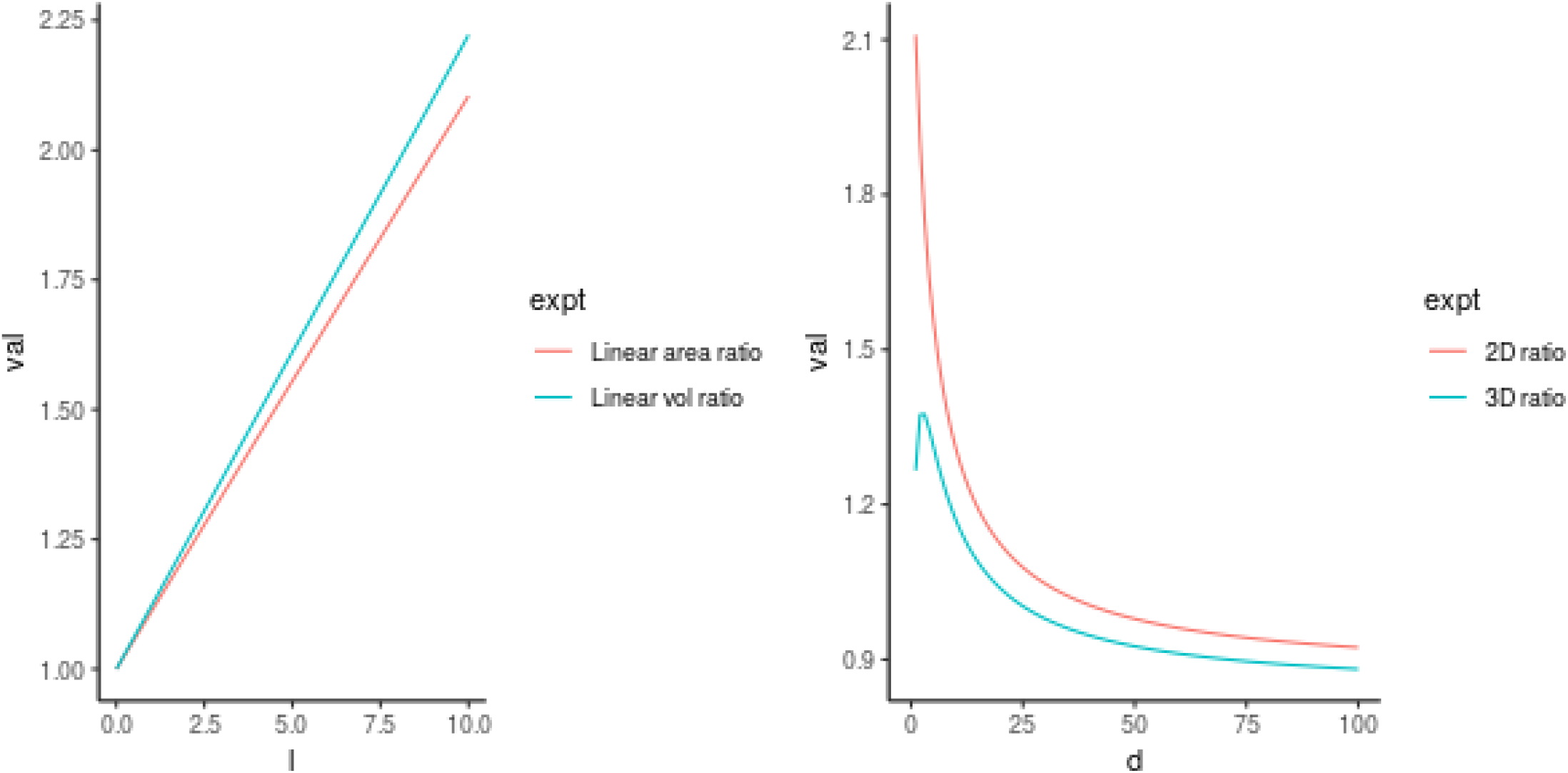
Advantages of stromules. (A) Increase of cytoplasmic area within a distance of 1 *µ*m of the plastid as stromule length *l* increases, plotted relative to the area without a stromule. Lines show the cases when membrane area or internal volume are conserved. (B) Size of cytoplasmic region within a distance *d* of the plastid when a 10*µ*m stromule is extended, relative to the no-stromule case. Relative increase is greater in 2D (area) than in 3D (volume).

### Simplified geometry in 2D vs 3D

Given that the stromule radius is rather smaller than the other length scales, we can simplify this picture further. We consider the *ρ* → 0 limit, where the stromule is a line with negligible width. We picture the body of the plastid as a circle, then compute the area of a shell around the plastid within distance *d*:

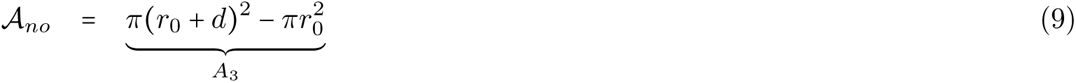

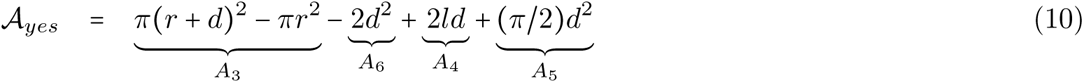

In 3D we can picture the body of the plastid as a cylinder with radius *r*_0_ and height *h* (Fig. 3C-D). The volume of a shall around the plastid within distance *d* is

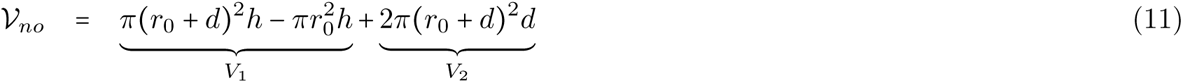

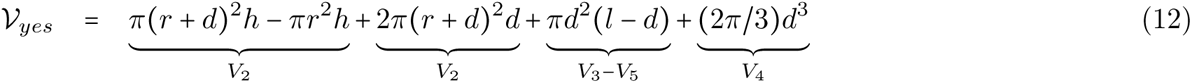

Fig. 4B shows the relative difference in cytoplasmic region size within distance *d* of the plastid in the yes-stromule and no-stromule cases. In 2D, the relative increase in area is always higher than the relative increase in volume in 3D, especially for low *d*.

### Branching angles

When a stromule changes angle or branches, the shell regions surrounding different segments of its structure may overlap. Fig. 3E shows an illustration of this, where an acute angle between stromule segments leads to the width-*d* shells of the two segments overlapping in the highlighted region. This overlap ‘double counts’ the corresponding region, so that overall less space is contained within a distance *d* of the plastid.

To compute how much space is lost in this way, we first compute the area of the highlighted region in Fig. 3E as a function of *α*, the angle between the two stromule segments.

Triangle *OAB* has base *AB* and height *d* sin *γ*. Triangle *ABC* has base *AB* and height 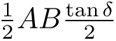. Quadrilateral *OACB* thus has area 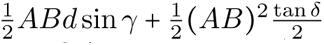.

Circle sector ⌔*OAB* has area *πd^2^ β*/(2*π*) = *βd^2^* /2. So the region of interest has area

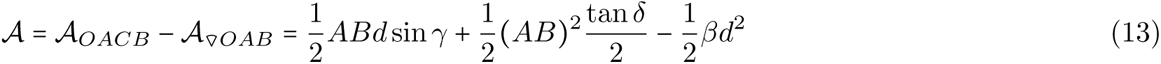

As length *OA* = *d*, *AB*/sin *β = d/*sin *γ*, for AB = d sin *β*/ sin *γ*.

As angles ∠*OAC = d,* and ∠*OBC* are right angles, *β* = *π* − *α*. By internal angles of triangles, *γ* = (*π* − *β*)/2 = *α*/2, *δ* = (*π* − *α*)/2.

Hence AB = *d* sin *α*/ sin *α*/2 and overall

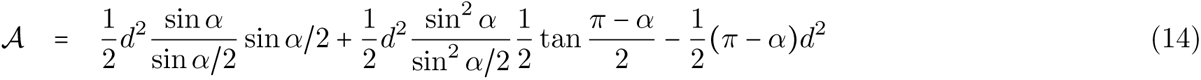

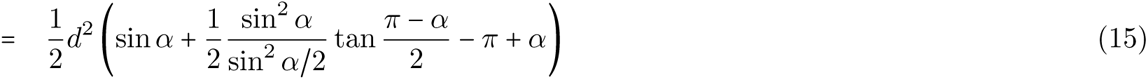

Now in the case of a branch point with three stromule segments *P, Q, R* incident on a branch point, there are two cases that require consideration. Without loss of generality, label *P* and *Q* as the segments with the largest angular separation. Then *R* can either fall ‘between’ or ‘outside’ the angle formed by *P* and *Q* (Figs. 3F and 3G respectively). In the ‘between’ case, the total overlap area is *A_PR_* + *A_QR_* − *A_PQ_*, where the overlap of overlaps is subtracted to avoid double counting. In the ‘outside’ case, the total overlap area is the sum *A_PR_* + *A_QR_* + *A_PQ_*. Then, treating the incident angle of *P* as the zero reference angle, we have

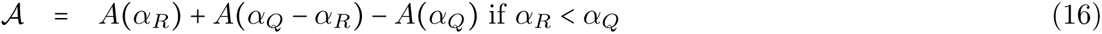

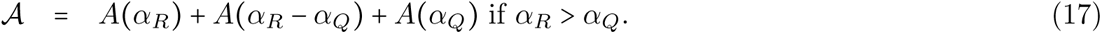

Fig. 5 shows this overall overlap area as a function of the two angles, with a minimum at *α_Q_* = 2*π* 3 and *α_R_* = 4*π* 3 (120^○^ separation).

---

**Figure 5:**
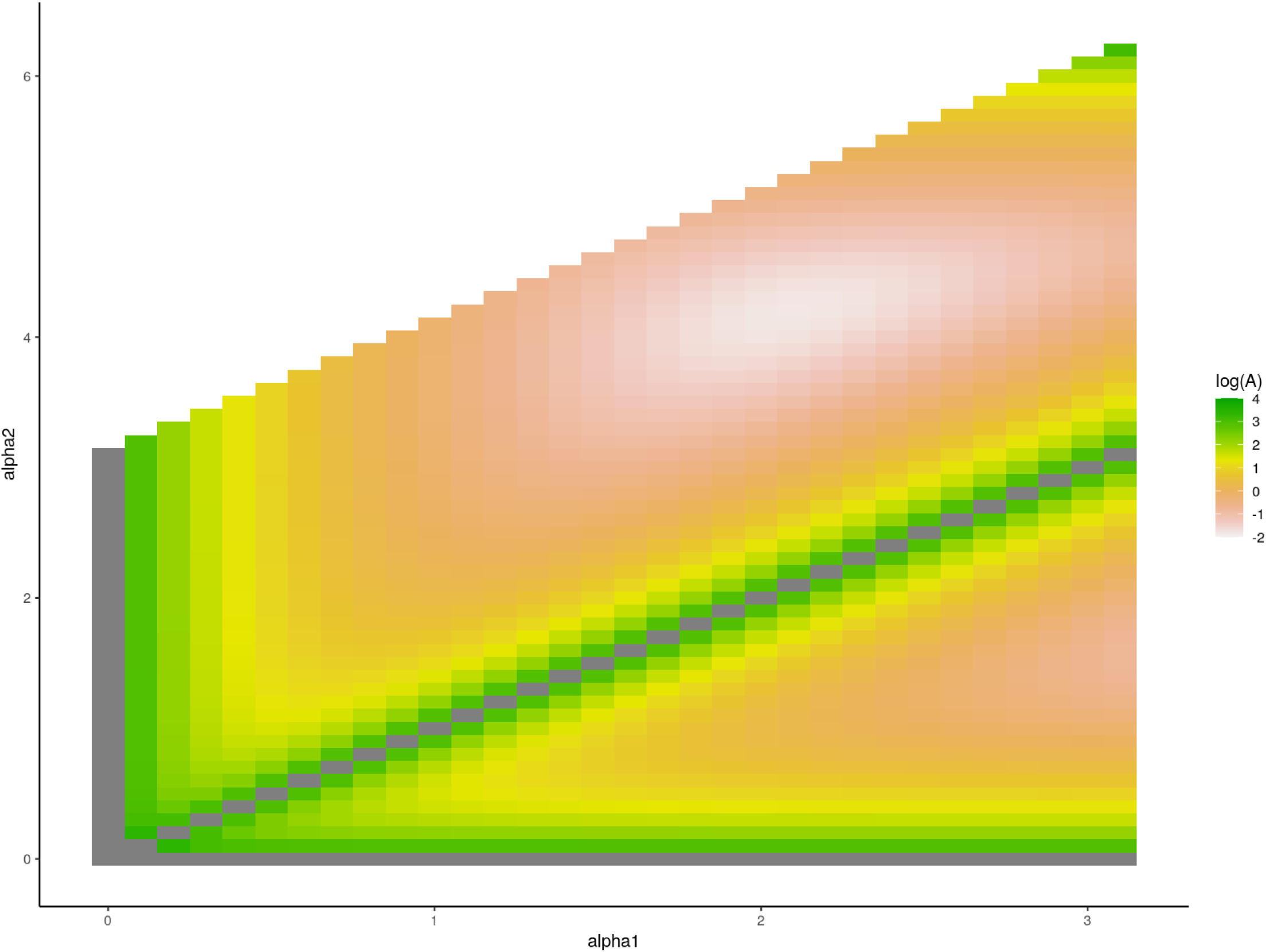
Optimal branching angles. Overlap area as a function of angles *α*_1_ and *α*_2_ in a stromule junction (logarithmic scale). Overlap area is minimised when angles are 2π*/*3 ≃ 2.1 and 4π*/*3 ≃ 4.2. A secondary minimum occurs at angles π ≃ 3.1 and π*/*2 ≃ 1.6. When angles are equal, or either are zero, stromule segments overlap perfectly and the overlap area diverges. Given the symmetry of the system, results are plotted only for *α*_1_ < π and *α*_2_ < *α*_1_ + *π* (values outside these inequalities can be mapped to this plot by relabelling the segments and angles).

**Figure 6:**
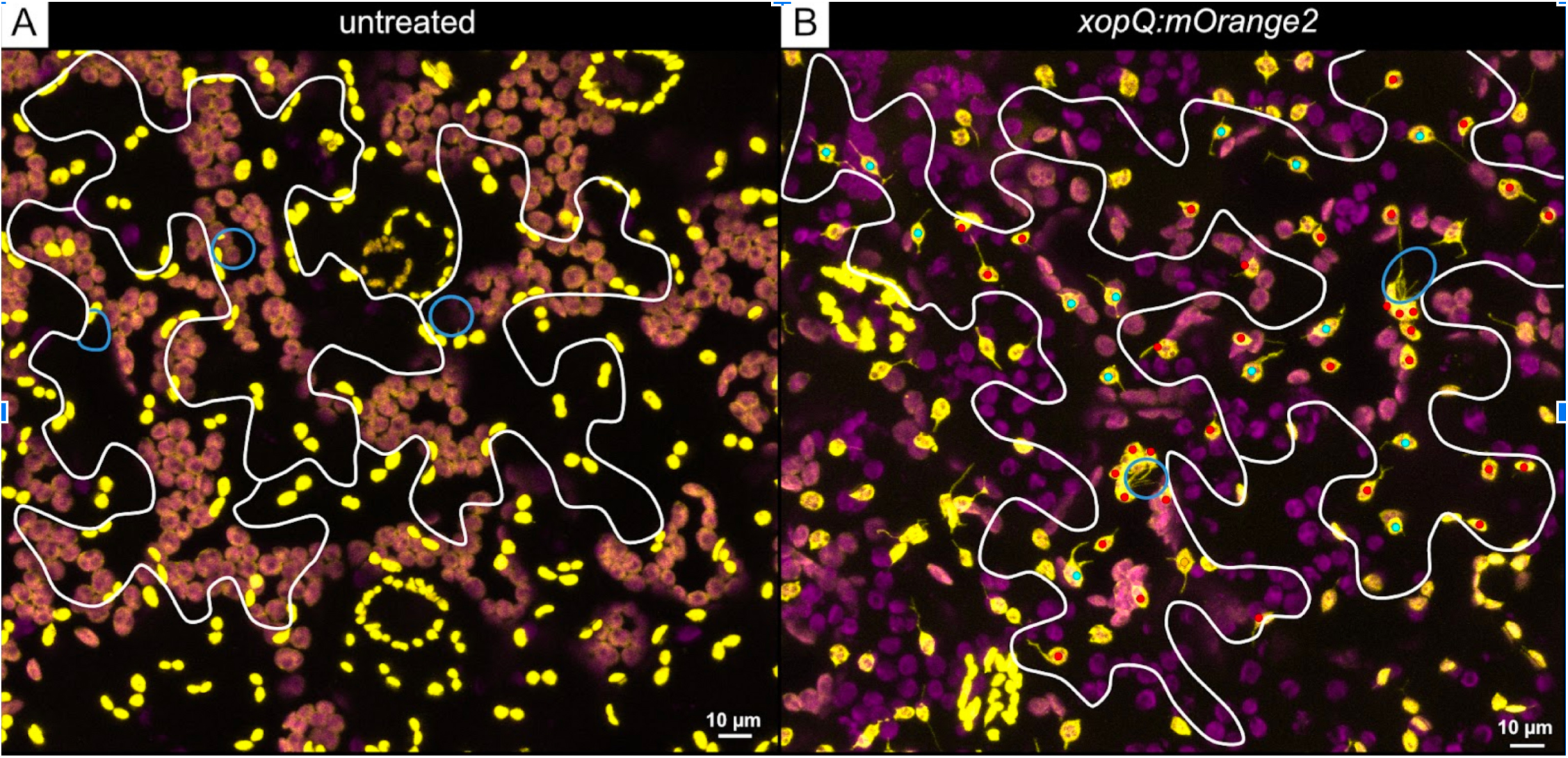
Maximum intensity projection along the z-axis of CLSM images capturing the lower epidermis cells of untreated (A) and treated (B) leaf sectors of a *N. benthamiana* transgenic line FNR:eGFP#7-25. The treated area was inoculated with *A. tumefaciens* cells mediating the expression of xopQ:mOrange2. White lines = cell outlines of pavement epidermis cells; blue lines = indicates the localisation of nuclei; coloured dots indicate plastids with stromules, red = 1, cyan = 2, yellow =

